# Genome-wide identification of MAPK gene family and comparative transcriptional profiling among different organ and stress response in two jute species

**DOI:** 10.1101/2024.04.02.587793

**Authors:** Borhan Ahmed, Anika Tabassum, Kazi Khayrul Bashar, MW Ullah, Nasima Aktar, MS Roni, Fakhrul Hasan, Mobashwer Alam

## Abstract

Mitogen activated protein kinase (MAPK) cascade is evolutionary conserved universal signal transduction module that plays central role in the growth and development of plants as well as in biotic and abiotic stress response. Although, MAPKs have been investigated in several model plants, no systematic analysis has been conducted in jute species (*Corchorus olitorius* and *C. capsularis*) even though, their genome sequencing has been completed. In the present study we identified 11 and 12 putative MAPKs in *C. olitorius* and *C. capsularis* using their genomic database, respectively. Here we provide a comprehensive bioinformatics analysis of the MAPK family from both *Corchorus* species including identification and nomenclature, chromosomal localization, sequence alignment, domain and Motif, gene structure, phylogenetic, functional analysis and investigation of expression analysis in response to abiotic stress and fiber cell development. The phylogenetic analysis of predicted MAPKs were clustered into four different clades and assigned with specific name based on their orthology based evolutionary relationship with *Arabidopsis*. Structural analysis of the MAPK genes revealed that there was a large variation among the exon number in both *Corchorus* species ranged from 2 to 11 but genes with the same clade had similar exon-intron structure. The sequence alignment analysis concede the presence of several conserved domain and motif including crucial signature phosphorylation motif TDY or TEY where first one is harbor in group D sequence and rest of the sequence contain TEY motif in their activation loop. Transcriptome analysis against salinity, drought along with fiber cell formation showed that MAPK4-1 genes in both jute genome highly expressed and may play a potential role in jute on adverse condition as well as jute fiber formation. These findings yielded new insights into the transcriptional control of MAPK gene expression, provide an improved understanding of abiotic stress responses and signaling transduction in jute, that lead to potential applications in the genetic improvement of jute cultivars.

## Introduction

Protein kinases (PK), the most prominent superfamily in the elaborate matrix of signal transduction proteins, allow cells of eukaryotic organisms to grow and development in a coordinated manner (Nolen et al., 2004; Tena et al., 2011; Zulawski et al., 2014). Mitogen-activated protein kinase (MAPK) has formed a PK member, which is evolutionarily conserved and plays role in transducing extracellular stimuli into intracellular responses in eukaryotes (Xu and Zhang, 2015). The signal transduction modules of MAPK play a significant role in the plant’s growth, development, and regulation of various abiotic and biotic stresses, such as drought, low temperature, high salt, mechanical damage, osmotic stress, oxidative stress and pathogen infection (Cardinale et al., 2002; Colcombet and Hirt, 2008; Jonak et al., 2002; Sinha et al., 2011). The MAPKs are important signaling molecules to perceive various signals and transduces them for active responses. These compounds carryout diverse phosphorylation processes at the transcriptional, translational and post-translational level by catalyzing the addition of phosphate groups to serine and threonine/tyrosine residues in their target proteins in both prokaryotic and eukaryotic cells (Ellis, 2012; Gonzalez Besteiro and Ulm, 2013). These modifications have led to changes in catalytic activity, affinity and interaction activity of target protein. However, the phosphorylation events in proteins are reversible due to protein phosphatase, enabling maintenance of the balance between kinase driven phosphorylation and phosphatase driven dephosphorylation events (Gonzalez Besteiro and Ulm, 2013).

The MAPK cascades are composed of three protein kinase modules: MAPK kinase kinase (MAPKKKs/MEKKs), MAPK kinase (MAPKKs/MKKs) and MAPK (MAPKs/MPKs)(Nishihama et al., 1995). MAPKs are activated when both tyrosine and threonine residues in the TEY/TDY motif are phosphorylated by dual-specificity kinases, MAPKKs, which in turn are activated by MAPKKKs via phosphorylation on two serine/threonine residues in the conserved Thr/Ser motif (Zhang et al., 2006).

MAPKs contains eleven domains (I–XI) that are found in all serine/threonine protein kinases. Threonine and tyrosine residues exist in between well conserved domains VII and VIII. Phosphorylation is required for the activation of the MAPKs (Hirt, 1997). The high degree of sequence similarity of MAPKs might reflect their functional conservation among different plant species. In all plant species, MAPKs carry either a Thr-Glu-Tyr (TEY) or Thr-AspTyr (TDY) phosphorylation motif at the active site. In contrast to TEY MAPKs, all TDY MAPKs have long C-terminal extensions. To date, functional data for TDY MAPKs is only available for one rice MAPK, BWMK1 (blast- and wound-induced MAP kinase). By contrast, the TEY MAPKs have been studied in many plant species including Arabidopsis, Medicago, tobacco, tomato, parsely and rice (Cheong et al., 2003). MAPKs regulate diverse cellular programs by relaying extracellular signals to intracellular responses (Cargnello and Roux, 2011). Plant MAPKs play an important role in the response to a broad variety of biotic and abiotic stresses including pathogen infection, wounding, temperature, drought, salinity and plant hormones regulations (Lee et al., 2011; Romeis, 2001; Tena et al., 2001; Zhang and Klessig, 2001). The Arabidopsis contains 20 MAPK genes and MPK3, MPK4 and MPK6 are mainly involved in stress responses (Ichimura et al., 2002; Jonak et al., 2002). In tobacco, ozone induces the activation of a p46 MAPK within minutes (Samuel et al., 2000). It has been proven that the activation of salicylate induced protein kinase (SIPK) and major ROS-induced MAPK, was important for the effective control of ROS damage (Samuel and Ellis, 2002). Although the ozone-induced activation of MPK3 and MPK6 in Arabidopsis was independent of ethylene, it was dependent on salicylic acid and resulted in the nuclear translocation of MAPKs (Ahlfors et al., 2004). Several MAPKs have been shown to be activated by osmotic stresses or NaCl in Medicago and tobacco (Jonak et al., 2002). MPK4 and MPK6 in Arabidopsis were phosphorylated and activated by MKK2 which was specifically activated by cold and salt stress (Teige et al., 2004). GhMPK6 maintains a high phosphorylation level in elongating fibers, and its phosphorylation was enhanced in fibers by phytohormones brassinosteroid (BR), ethylene and indole-3-acetic acid (IAA)(Chen et al., 2020).

Advancements in sequencing technologies and bioinformatics tools have greatly increased the pace of genome sequencing projects, resulting in successful sequencing of several plant genomes. Post genome sequencing projects have enabled relatively easy identification of particular gene families based on conserved signature motifs and sequence similarity. Available genome sequences from several plants have provided us with an opportunity to identify MAPK family members across photosynthetic eukaryotes (plants and algae) that shed light on MAPK evolution and signaling in plants and lower photosynthetic eukaryotes (Ichimura et al., 2002).

Considering the importance of biodegradable natural fiber production and environment protection as well as the emergence of MPPK genes as promising candidate for plant stress tolerance modification, it was of interest for us to characterize the MAPK gene family in two genome decoded cultivated jute species.

Jute is an agricultural commodity that singly earns foreign exchange equivalent to 38.69 billion taka annually to our national economy, which is the second largest source of foreign exchange (Islam et al., 2017). Due to the national food demand is increasing sharply; as a result jute is being pushed to the marginal lands. Jute is very sensitive to abiotic stresses like salinity, drought, water logging, low temperature and short day length. Among these abiotic stresses, salinity and drought are the most severe environmental stress that constraints global crop production including Bangladesh. Since jute is a bast fiber bearing crop, so our aim is also to identify the effective MAPK signaling gene(s) that enhance fiber quality as well as productivity. Furthermore, genome-wide analysis of MAPK family can provide a strong groundwork for the identification of candidate salt, drought stress tolerance as well as potential fiber formation related gene(s) in jute along-with their expression with signaling regulation.

## Materials and method

### Genomic Identification of putative MAPK genes

To identify the MAPK gene sequences in *Corchorus olitorius* and *C. capsularis*, their protein and genome sequence were downloaded from Basic and Applied Research on Jute Project (BARJ) database. The MAPK protein sequence of *Arabidopsis thaliana* available in TAIR database were used as query sequence to execute Blastp and tBlastn algorithm program with cut off e-value of e^-10^ against protein and genome sequence of both *Corchorus* species genome to identify the putative MAPK genes. The identified MAPK proteins of both jute genome were further screened to confirm the presence of serine/threonine protein kinase domain (PF00069) used by Pfam (https://pfam.xfam.org/) along with signature motif TXY (TDY or TEY) in activation loop were verified manually through multiple sequence alignment.

### Physico molecular characterization of MAPKs

The physico-chemical analysis of MAPK genes was performed using online ExPasy tool ProtParam (http://expasy.org/tools/protparam.html)(Gray, 1996) to determine the length of protein, molecular weight (MW), theoretical values of isoelectric point (pI), the grand average of hydropathicity (GRAVY) and other related properties. In order to predict the subcellular localization of Co and CcMAPKs, Plant-mPloc server (http://www.csbio.sjtu.edu.cn/bioinf/plant-multi/)(Chou and Shen, 2010) was used.

### Sequence alignment, phylogenetic analysis and naming of MAPKs

The multiple sequence alignments of the amino acid sequence of MAPKs of both *Corchorus* spp along with *AtMAPK6* were conducted using clustal omega(Sievers and Higgins, 2018) with the default settings. The alignment data were further processed using ESPript (http://espript.ibcp.fr) with default parameter except number of sequence per column 100. In order to understand the relationship among the MAPK genes in arabidopsis, cacao and both *Corchorus* species, a phylogenetic tree was constructed by MEGA7 software(Kumar et al., 2016). Initially, multiple sequence alignment of abovementioned 4 species MAPK protein sequences were created using ClustalW tool in MEGA7 and then according to alignment file, a phylogenetic tree was generated using the neighbor-joining (NJ) method inferred from 1000 bootstrap replicates with other default parameter. The naming of *Co* and *CcMAPKs* genes were assigned according to phylogenetic tree showing orthology with Arabidopsis and Cacao along-with their reciprocal BLASTP identity.

### Chromosomal location, structural and motif analysis

For Chromosomal distribution and gene structure analysis the genomics DNA sequences, chromosome number, start and end base pairs, Generic Feature Format (gff3) data of *C. olitorius* and *C. capsularis* were retrieved from BARJ databse. The physical location of *Co and CcMAPKs* on each chromosome/scaffold was detected using BLASTNT search against the genome database of the *C. olitorius* and *C. capsularis* respectively. Start sites of the MAPKs genes were used as the indicative position of the genes on the chromosome or the scaffold. Due to the unavailability of the full chromosome-scale assembly in both jute genome, led also to use assembled sequence in the form of scaffolds to determine the position of *Co* and *CcMAPKs*. The position and structures of exons and introns along with the corresponding phylogenetic tree were illustrate of MAPK genes in both jute genomes were determined by online tools Gene Structure Display Server, GSDS(Hu et al., 2015) (http://gsds.cbi.pku.edu.cn/) using *Co* and *CcMAPK* annotation file in Generic Feature Format Version 3 (gff3) data. Conserved motifs of the identified gene family in both jute genome were predicted using MEME (Multiple expectation maximization for motif elicitation) discovery server (http://meme.sdsc.edu/meme/meme.html)(Bailey et al., 2009) with default settings excepts maximum number of motif 15.

### Promoter and functional analysis of identified MAPK genes

Gene Ontology (GO) annotation of *Co* and *CcMAPKKs* were conducted for describing the biological processes, cellular components and molecular functions were determined by Blast2GO program(Gotz et al., 2011) using amino acid sequences with default parameters. To identify putative *cis*-acting regulatory elements in the promoter sequences of the identified MAPKK family genes, 1Kbp upstream intergenic region from the initiation codon (ATG) of the predicted transcription sites were extracted from jute genome data (www.jutegenome.org). The PlantCARE (http://bioinformatics.psb.ugent.be/webtools/plantcare/html/)(Rombauts et al., 1999) and PLACE (http://www.dna.affrc.go.jp/PLACE/)(Higo et al., 1999) databases were used to confirm the putative *cis*-elements in the promoters.

### Expression Analysis

For the expression pattern analysis of *Cc* and *CoMAPK* genes under drought and salinity stresses, & seedling and fiber cell publicly available transcriptome data were downloaded from Sequence Read Archive (SRA) (https://www.ncbi.nlm.nih.gov/sra). The accession number and sample information of the used data were listed in Table S1. Gene expression under salinity stress was explored using the available data from two salt-tolerant varieties Yueyuan No.5 (YY) (C. *capsularis*)(Yang et al., 2017c) and TC (C. *olitorius*)(Yang et al., 2017b), and a salt-sensitive accession NY/253C (NY)(Yang et al., 2017b). For drought stress study, data from a drought-tolerant *C. olitorius* (Gangfengchangguo, GF)(Yang et al., 2017a), and a drought-sensitive *C. capsularis* (Yueyuan No.5, YY)(Yang et al., 2017a) were used. To explore the expression pattern for fiber development, we used the transcriptomic data of seedling and fiber of both species (Accession id: SRX2369402, SRX2369404, SRX2369401, SRX2369403)(Islam et al., 2017).

All the transcriptome sequencing data obtained from Illumina sequencing were quality checked by FastQC v.0.11.9 (Andrews S 2010) The raw data were filtered to trim low-quality reads by Trimmomatic *v.*0.36(Neupane et al., 2019). High-quality clean RNA-Seq reads were mapped to both *Corchoru*s species data by TopHat2 (version 2.1.0, Baltimore, MD, USA) with the default settings(Kim et al., 2013). The mapped reads were fed into Cufflinks2 v.2.1.1 suite(Trapnell et al., 2012) for transcriptome assembly and differential expression analysis. Then Cuffdiff2 calculated the *p-values* for differentially expressed genes (DGE), based on the normalized Fragments Per Kilobase Of Exon Per Million Fragments Mapped (FPKM) values. The clustered heatmap of Z scaled FPKM values of Co and CcMAPKs was generated using the heatmap function of R package (version 3.2.2; available online: https://cran.r-project.org/web/packages/pheatmap/).

## Results

### Genome wide identification of MAPK gene family and their physico-chemical properties

The query of 20 MAPK genes of *A. thaliana* through the BLAST searching, we have identified a total of 11 and 12 putative MAPKs in *C. olitorius* and *C. capsularis* genome respectively. All the identified *Co* and *CcMAPKs* were further validated by their characteristics serine/threonine protein kinase domain (PF00069) using Pfam tool and manually analyzed the presence of signature motif (TEY or TDY) using sequence alignment (Hamel et al., 2006) (Fig. 1). Genes were named following the MAPK genes nomenclature guideline (Hamel et al., 2006; Ichimura et al., 2002) for plants, a two letter prefix derived from the genus and species names of the organisms in which the genes are present. *Co* and *Cc* prefixes were used for *Corchorus olitorius* and *Corchorus capsularis*, respectively and the rest of the part of the genes were named following the Arabidopsis and Cacao orthologs of phylogenetically related MAPK genes of same clade (Johanson and Gustavsson, 2002) then number was followed by a hyphen with a number if paralogs were present. Such guidelines for nomenclature of MAPKs have been employed in previous studies (Chen et al., 2012; Goyal et al., 2018; Hamel et al., 2006; Liang et al., 2013; Liu et al., 2015; Mohanta et al., 2015; Neupane et al., 2013; Piao et al., 2018; Wang et al., 2017). The numbers of amino acids in the MAPK sequences were ranged from 366 (*CoMAPK13*) - 612 (*CoMAPK20*) in *CoMAPKs* and 366 (*CcMAPK7*) - 609 (*CcMAPK20*) in *CcMAPKs*. The molecular weight of *Co* and *CcMAPKs* ranged from 69.91 to 42.18 and 69.69 to 42.44 kDa, respectively. Like their orthologs the isoelectric point of MAPKs proteins were slightly acidic to moderately alkaline in nature (5.27 – 9.28). The negative GRAVY value indicating their hydrophobic nature. All the MAPKs genes in both jute genomes were localized in nucleus according to Plant-mPLoc server.

**Figure 1.**
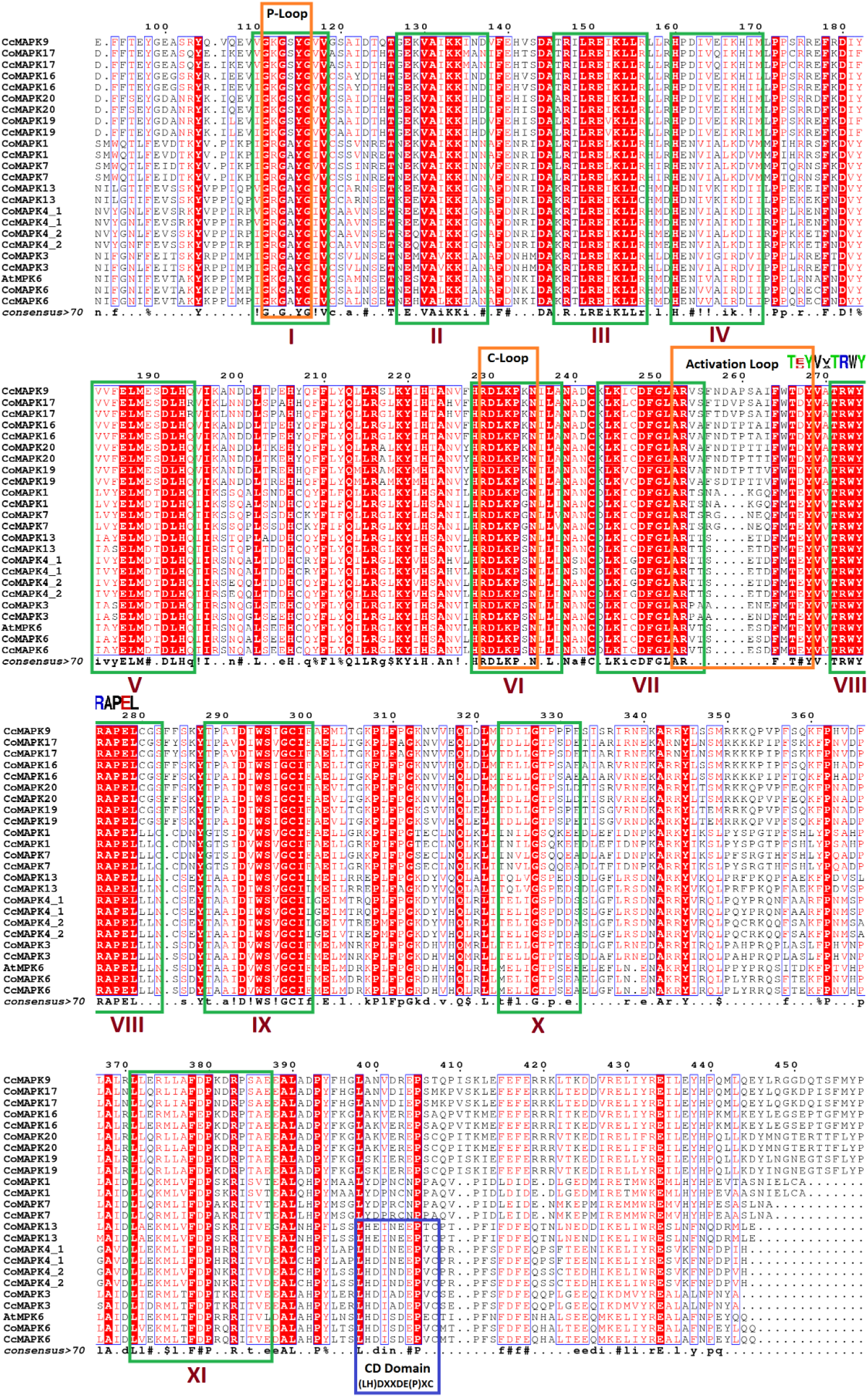
Multiple sequence alignment of MAPK genes in *C. olitorius* and *C. capsularis.* The figure shows presence of I-XI conserve domain, P-loop, C-loop, Activation loop and CD domain with proper marking

### Sequence alignment and identification of conserved domain

Multiple sequence alignment of all the identified *Co* and *CcMAPKs* along with well characterized AtMAPK6 amino acid residues was conducted (Fig. 1). All of these MAPKs contained the 11 conserved kinase subdomains and a TXY motif (T[D/E]YV[V/A]ATRWYRAPEL) in the activation loop between the kinase catalytic subdomains VII and VIII. Furthermore, the N-terminal regions of the 23 *Co* and *CcMAPKs* have a glycine-rich motif (GRG[A/S]YG) located in the phosphate-binding loop (P-loop) in subdomain I, which acts as an ATP- and GTP-binding site (Saraste et al., 1990) (Fig 1). Sequence comparison of the conserved amino acid motif TXY, which is phosphorylated by MAPKKs, classified MAPKs into two subtypes: one is those containing the amino acid motif TEY (TEY subtype) and another is those with the amino acid motif TDY (TDY subtype) at the phosphorylation site. The TEY subtype found in three Groups, A, B and C, whereas the TDY subtype forms a more distant Group D in accordance with phylogeny (Fig. 1).

**Table 8.3.2.** Physico-chemical properties of MAPK gene family in *C. olitorius* (a) and *C. capsularis* (b)

| Gene Name | Protein ID | Protein length | Theoretical pI | Molecular Formula | Aliphatic index | Instability index | GRAVY | Molecular weight (Da) |
| --- | --- | --- | --- | --- | --- | --- | --- | --- |
| 1 | CcMAPK1 | 372 | 6.53 | C<sub>1924</sub>H<sub>3013</sub>N<sub>513</sub>O<sub>544</sub>S<sub>21</sub> | 96.75 | 40.73 | -0.224 | 42708.46 |
| 2 | CcMAPK7 | 368 | 8.27 | C<sub>1906</sub>H<sub>3006</sub>N<sub>518</sub>O<sub>535</sub>S<sub>22</sub> | 95.38 | 35.63 | -0.240 | 42443.30 |
| 3 | CcMAPK4_2 | 374 | 6.20 | C<sub>1912</sub>H<sub>2991</sub>N<sub>525</sub>O<sub>564</sub>S<sub>16</sub> | 88.34 | 42.11 | -0.358 | 42869.92 |
| 4 | CcMAPK4_1 | 374 | 6.28 | C<sub>1929</sub>H<sub>3005</sub>N<sub>537</sub>O<sub>558</sub>S<sub>14</sub> | 90.96 | 42.20 | -0.357 | 43096.18 |
| 5 | CcMAPK6 | 400 | 5.39 | C<sub>2056</sub>H<sub>3193</sub>N<sub>557</sub>O<sub>604</sub>S<sub>20</sub> | 87.10 | 43.71 | -0.371 | 46019.54 |
| 6 | CcMAPK3 | 375 | 5.59 | C<sub>1916</sub>H<sub>2977</sub>N<sub>525</sub>O<sub>555</sub>S<sub>17</sub> | 92.37 | 38.27 | -0.265 | 42791.92 |
| 7 | CcMAPK13 | 371 | 5.27 | C<sub>1916</sub>H<sub>3009</sub>N<sub>509</sub>O<sub>566</sub>S<sub>18</sub> | 93.32 | 44.26 | -0.303 | 42808.12 |
| 8 | CcMAPK16 | 563 | 8.80 | C<sub>2870</sub>H<sub>4508</sub>N<sub>796</sub>O<sub>830</sub>S<sub>23</sub> | 77.99 | 36.78 | -0.475 | 64181.58 |
| 9 | CcMAPK20 | 609 | 9.08 | C<sub>3126</sub>H<sub>4913</sub>N<sub>867</sub>O<sub>899</sub>S<sub>21</sub> | 80.08 | 41.28 | -0.467 | 69698.93 |
| 10 | CcMAPK9 | 599 | 7.06 | C<sub>3026</sub>H<sub>4744</sub>N<sub>854</sub>O<sub>904</sub>S<sub>15</sub> | 78.98 | 43.90 | -0.538 | 68033.04 |
| 11 | CcMAPK19 | 594 | 9.28 | C<sub>3022</sub>H<sub>4756</sub>N<sub>840</sub>O<sub>862</sub>S<sub>17</sub> | 82.91 | 38.00 | -0.407 | 67193.14 |
| 12 | CcMAPK17 | 550 | 8.47 | C<sub>2808</sub>H<sub>4440</sub>N<sub>784</sub>O<sub>816</sub>S<sub>23</sub> | 85.44 | 42.37 | -0.401 | 62976.29 |

**b.**
| Gene Name | Protein ID | Protein length | Theoretical pI | Molecular Formula | Aliphatic index | Instability index | GRAVY | Molecular Weight (Da) |
| --- | --- | --- | --- | --- | --- | --- | --- | --- |
| 1 | CoMAPK1 | 372 | 6.53 | C<sub>1925</sub>H<sub>3015</sub>N<sub>513</sub>O<sub>545</sub>S<sub>21</sub> | 96.48 | 40.50 | -0.230 | 42738.48 |
| 2 | CoMAPK7 | 368 | 8.28 | C<sub>1905</sub>H<sub>3004</sub>N<sub>518</sub>O<sub>534</sub>S<sub>22</sub> | 95.38 | 36.34 | -0.229 | 42413.28 |
| 3 | CoMAPK4_2 | 374 | 6.20 | C<sub>1909</sub>H<sub>2993</sub>N<sub>525</sub>O<sub>564</sub>S<sub>16</sub> | 89.39 | 42.51 | -0.355 | 42835.90 |
| 4 | CoMAPK4_1 | 374 | 6.28 | C<sub>1930</sub>H<sub>3007</sub>N<sub>537</sub>O<sub>558</sub>S<sub>14</sub> | 90.96 | 41.80 | -0.357 | 43110.21 |
| 5 | CoMAPK3 | 375 | 5.54 | C<sub>1925</sub>H<sub>2983</sub>N<sub>525</sub>O<sub>555</sub>S<sub>17</sub> | 91.84 | 38.30 | -0.263 | 42906.07 |
| 6 | CoMAPK13 | 366 | 5.32 | C<sub>1899</sub>H<sub>2977</sub>N<sub>501</sub>O<sub>553</sub>S<sub>16</sub> | 97.51 | 42.66 | -0.240 | 42187.51 |
| 7 | CoMAPK16 | 558 | 8.65 | C<sub>2841</sub>H<sub>4455</sub>N<sub>785</sub>O<sub>822</sub>S<sub>22</sub> | 78.87 | 37.87 | -0.445 | 63465.71 |
| 8 | CoMAPK17 | 498 | 8.79 | C<sub>2550</sub>H<sub>4036</sub>N<sub>710</sub>O<sub>740</sub>S<sub>21</sub> | 82.83 | 41.11 | -0.446 | 57153.67 |
| 9 | CoMAPK20 | 612 | 9.12 | C<sub>3136</sub>H<sub>4934</sub>N<sub>870</sub>O<sub>901</sub>S<sub>21</sub> | 80.82 | 41.30 | -0.450 | 69914.22 |
| 10 | CoMAPK19 | 594 | 9.28 | C<sub>3022</sub>H<sub>4755</sub>N<sub>841</sub>O<sub>862</sub>S<sub>17</sub> | 82.74 | 38.15 | -0.414 | 67206.14 |
| 11 | CoMAPK6 | 400 | 5.34 | C<sub>2059</sub>H<sub>3197</sub>N<sub>557</sub>O<sub>606</sub>S<sub>20</sub> | 86.85 | 44.41 | -0.385 | 46091.60 |

The conserved catalytic loop (C-loop, HRDLKP[G/S/K]N), which harbors an Asp (D) residue that is the key site of the phosphorylation reaction, was also present in subdomain VI of identified MAPKs (Fig. 1). The MAPKs common docking (CD) domain consists of two negatively charged amino acid residues and functions as a docking site for MAPK kinases, substrates, and negative regulators (Tanoue et al., 2000). This conserved domain was observed in the extended C-terminal region of the *Co* and *CcMAPK* members belonging to the Group A and Group B (*Co* and *CcMAPK13, 4_1, 4_2, 3* and *6*), indicating that the acidic cluster (LH[D/E]XX[D/E]EPXC) may establish electrostatic interactions with upstream factors and form a modular recognition system that mediates connectivity (Fig. 1).

### Phylogenetic analysis of MAPK genes

A phylogenetic tree was constructed based on the protein sequences of 11 *CoMAPKs*, 12 *CcMAPKs*, 16 *TcMAPKs* and 20 *AtMAPKs* genes showed four distinct clusters representing different group of MAPKs (Fig. 2). Each pair of orthologous MAPKs genes has a high percentage of similarity in both nucleotide and amino acid sequence (>90% identity at the protein level). *Co* and *CcMAPKs* belonging to the A, B and C group, possess a TEY motif, whereas the group D possesses a TDY motif at the activation site. *Co* and *CcMAPK4_1, 4_2* and *13* genes are clustered in Group A, which contains well-characterized MAPK genes including *AtMPK4, 5, 11, 12,* and *13* genes. *Co* and *CcMAPK3* and *6* genes belong to Group B, which includes *AtMPK3, 6* and *10* genes. Group C contained two genes of each *Corchorus* species: *Co* and *CcMAPK1* and *7* that include *AtMPK1, AtMPK2, AtMPK7* and *AtMPK14*. Largest group D includes *Co* and *CcMAPK16, 17, 19, 20* and *CcMAPK9* along-with *AtMAPK8*, *15*, *16*, *17*, *19*, *20*.

**Figure 2.**
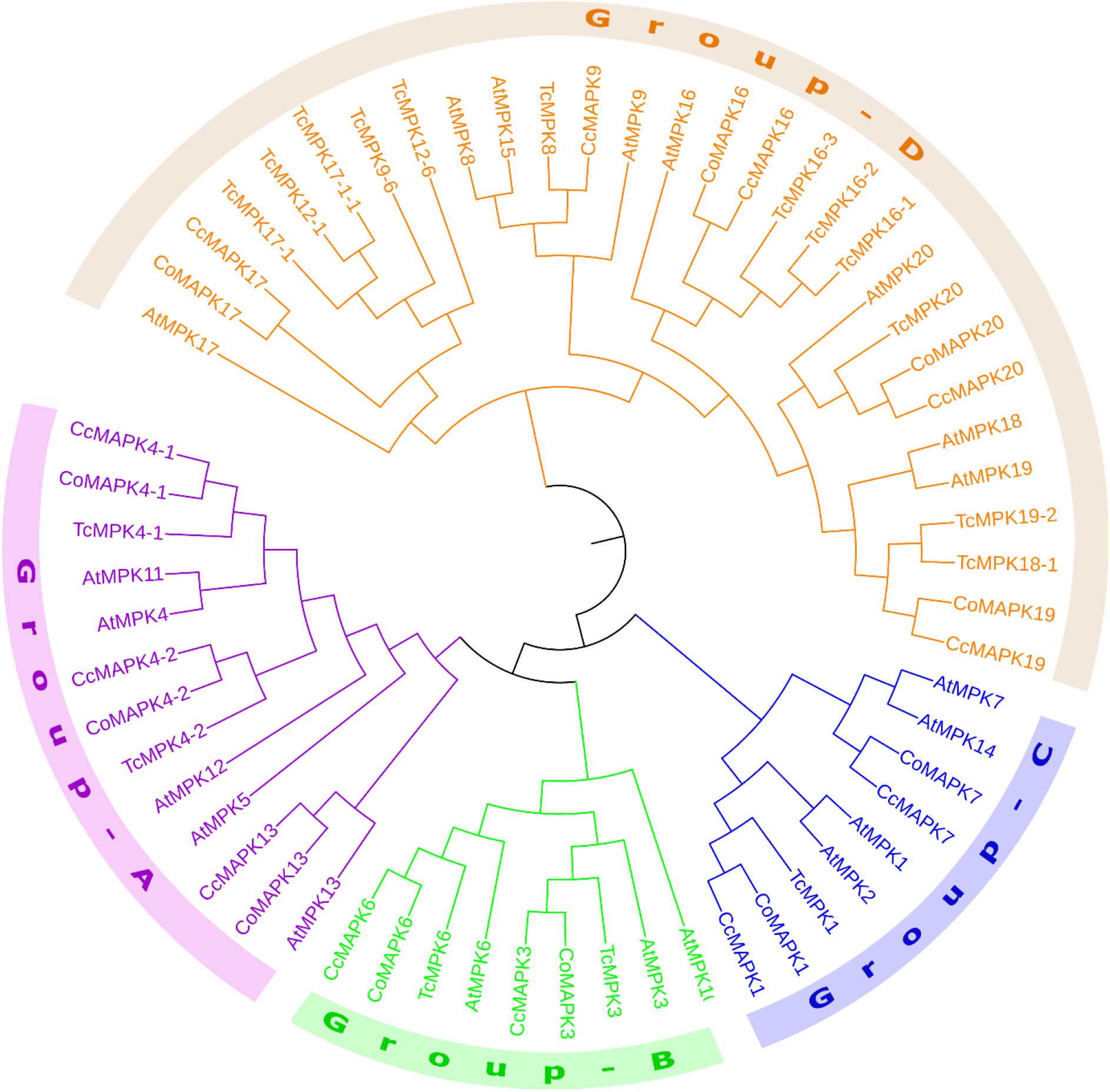
Phylogenetic relationship of putative *MAPK* genes in *C. olitorius, C. capsularis, A. thaliana* and *Theobroma cacao.* The phylogenetic tree was created using MEGA6.0 program with the neighbor-joining (NJ) method. Bootstrap values for 1000 replicates are indicated at each branch. Different colour indicate different groups of MAPKs.

### Chromosomal distribution, gene structure and conserved motif analysis

Among the 20 MAPKs, 7 and 8 MAPKs were identified in the chromosomes of *C. olitorius* and *C. capsularis*, respectively. The remaining MAPKs were identified on unanchored Scaffolds. The identified MAPK genes are distributed unevenly on the seven chromosomes in both *Corchorus* species with maximum number of genes (three MAPKs) on chromosome 1 in *C. capsularis* followed by 2 MAPKs in Chromosome 2 and 3 in both *Corchorus* species while none of the MAPK genes was found to be located on chromosome 6.

The identification of exon-intron structures for each *Co* and *CcMAPKs* gene were determined by aligning corresponding genomic DNA sequences. The exon/intron structures of putative *Co* and *CcMAPKs* genes could also be divided into four subgroups based on their phylogenetic relationship (Fig. 3). We found that *Co* and *CcMAPKs* genes in C and D groups have strikingly different exon/intron structures whereas group A and B consists same number of intron/exon but that the gene structures of putative *Co* and *CcMAPKs* members in the same group were highly conserved in both *Corchorus* species. (Fig. 3). The putative *Co* and *CcMAPKs* members were composed of six exons in both Group A and B whereas, Group C only had two exons. Nine to 11 exons were present in the Group D and a larger number of exons with variable exon lengths than other groups. From the figure 3 it was visible that exon size is more conserve among the group than intron length.

**Figure 3.**
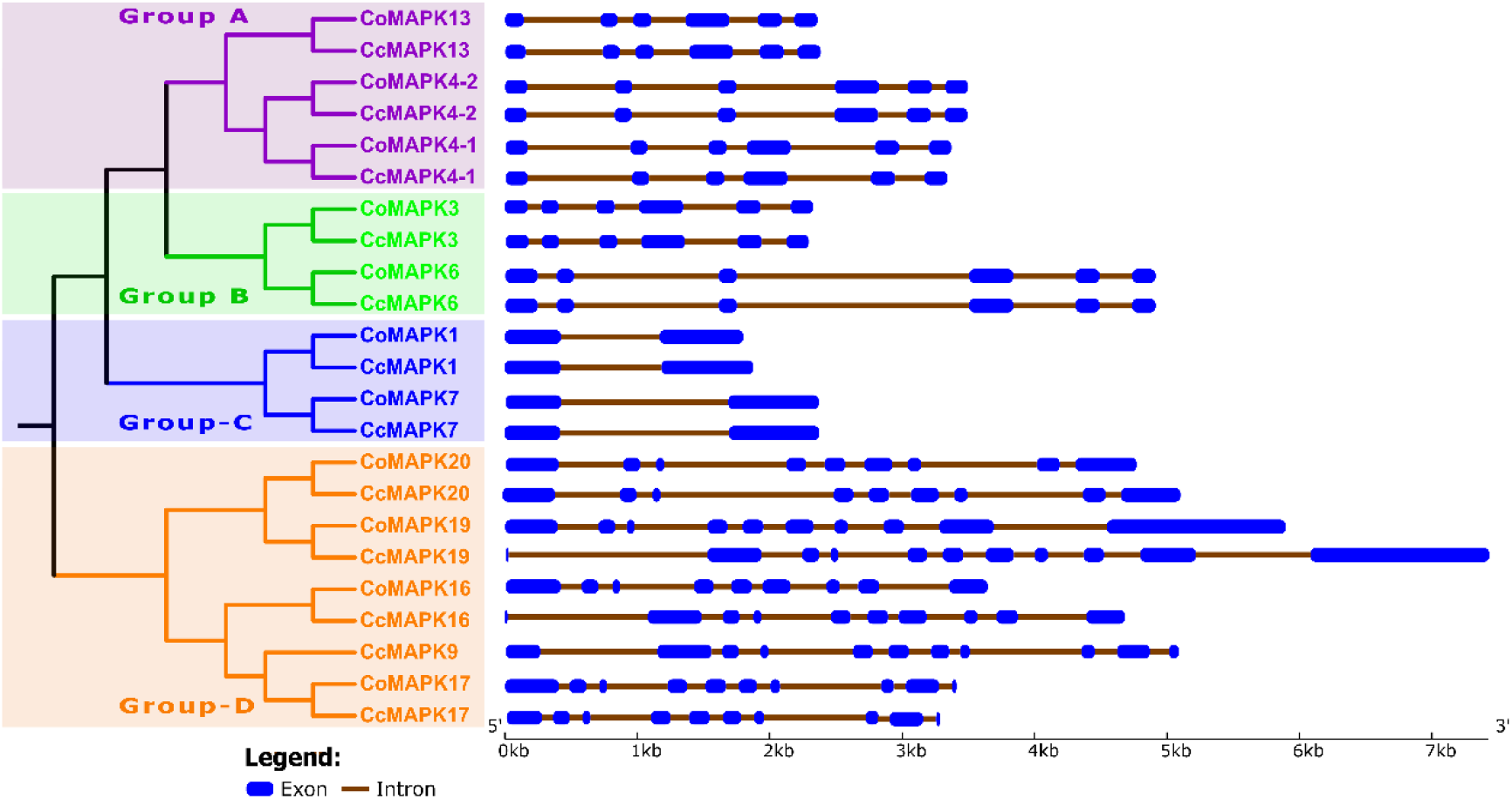
The phylogenetic analysis and intron/exon structures of putative *MAPK* genes in *C. olitorius* and *C. capsularis*. The phylogenetic tree (left panel) was created using MEGA6.0 program with the neighborjoining (NJ) method. Exon/intron structures of the *MAPK* genes are shown in the right panel. The yellow boxes indicate the exons, whereas the single lines indicate introns. Gene models were drawn to scale as indicated on bottom.

To explore the structural diversity of the *Co* and *CcMAPKs* genes, we submitted the 23 putative *Co* and *CcMAPKs* protein sequences to the online MEME program to search for conserved motifs (Figure 4). 15 conserved motifs were identified. Specifically, all the identified *Co* and *CcMAPKs* contained motifs 1 (Contained C-Loop), 2, 3 (P-Loop), 4 (CD Doamin), 5, 6 (Contained the TXY signature motif), 9, 12, and 13 (Fig. 4), indicating that all the *Co* and *CcMAPKs* were typical of the MAPK family. We found all the members identified in the same subgroup shared similar conserved motifs. For instance, along with all the conserved motifs, most MAPK proteins in Groups A and B had specific motif 10 and 15 but devoid of Motif 14 and 15 whereas Motif 11 is unique to group C but lack of motif 8, 10, 13, 14 and 15. In Group D Motif 14 is unique but absent of motif 8, 10 and 11.

**Figure 4.**
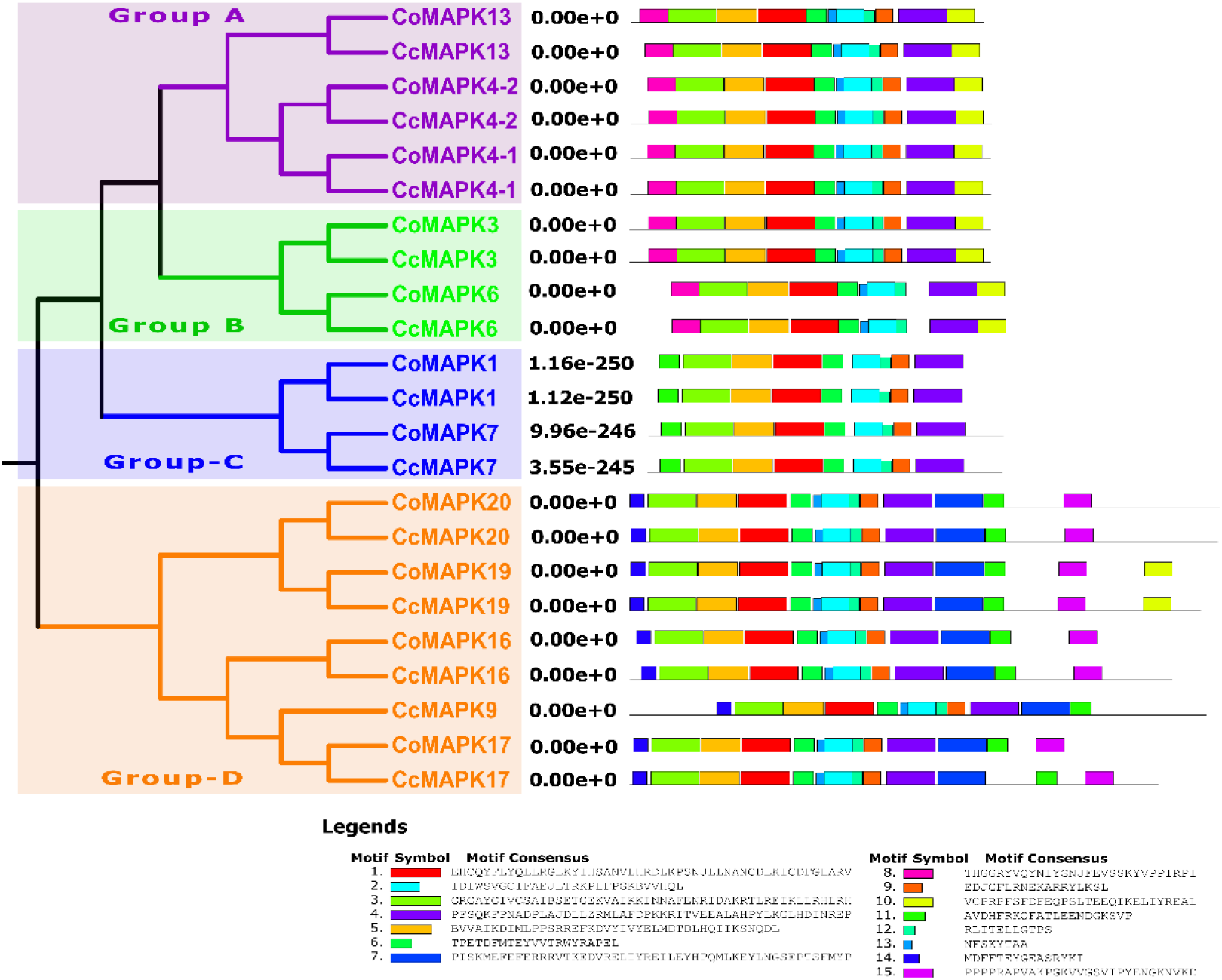
The conserved motifs of *C. olitorius* and *C. capsularis* putative MAPKs according to the phylogenetic relationship. All motifs were identified online with the MEME program with the complete amino acid sequences of the 11 CoMAPKs and 12CcMAPKs. Different colors of the boxes represent different motifs in the corresponding position of each Co and CcMAPK proteins.

### Gene ontology annotation and putative cis-element analysis of Co and Cc MAPKs genes

The Gene Ontology (GO) analysis performed using all the Co and CcMAPK proteins showed the putative participation in diverse biological process, molecular function and cellular components according to GO database. The analysis of biological process revealed that the MAPK signal transduction (MAPK Cascade GO:0000165) was predominant. Besides, identified MAPKs were involved in regulation of gene expression (GO:0010468), response to cold (GO:0009409) and other 22 different biological categories. The result also indicated that Co and CcMAPKs are mainly localized in nucleus (GO:0005634), and may also found in Phramoplast (GO:0009524), preprophase band (GO:0009574), trans-golgi network (GO:0005802) and integral component of membrane (GO:0016021). The molecular function clearly showed ATP binding and MAP Kinase were the main activities along with slightly RNA binding and tyrosine-tRNA ligase activity. In conclusion, functional analysis of *Co* and *CcMAPKs* suggest their involvement in signaling mechanism and kinase activity initiated from nuclear region.

Cis-elements modulate gene expression and provide an initial trigger for functional dissection of transcriptional sites among the upstream regions. To investigate the possible roles of MAPKs identified in the both jute genome, corresponding promoter regions (1 kb in length upstream region from the initiation codon ATG) of the *Co* and *CcMAPKs* genes were subjected to *cis*-element analysis by PlantCARE databse. (Fig. 5). Using the PlantCare database, we identified a total of 98 cis-element (four unnamed) in the promoter regions of jute. The identified cis-elements were divided into eight major groups, such as hormonal / environment responsive (ARE, AuxRR-core, ABRE, CGTCA-motif, TCA, WUN-motif etc.), Light responsive (GT1-motif, GATA-motif, G-box, GATA-motif, Box 4 etc.), site binding related element (Myb, MBS, CCAAT-box, MRE, AT-rich element etc.) and promoter core functional element (TATA-box, TATA, CAAT-box) (Fig. 5). The core promoter TATA-box and CAAT box had a greater number of promoter functions followed by unknown function and site binding related element. The well-known stress–response element (STRE, AAGGGG) and ABA responsible element (ABRE, C/TACGTGGC) was observed almost all the Co and CcMAPKS showing their response against multiple stress, including cold, drought, salt(Zhu et al., 2005),(2012). CGTCA-motif and the TGACG-motif participate in Methylejasmonate (MeJA) production the response to several environmental stresses. MeJA responsiveness also involved in multiple physiological processes, including plant growth and development, abscission, maturity, and secondary metabolism(Browse and Howe, 2008; Creelman and Mullet, 1997; Pieterse et al., 1998; Turner et al., 2002; Wang et al., 2011).

**Figure 5.**
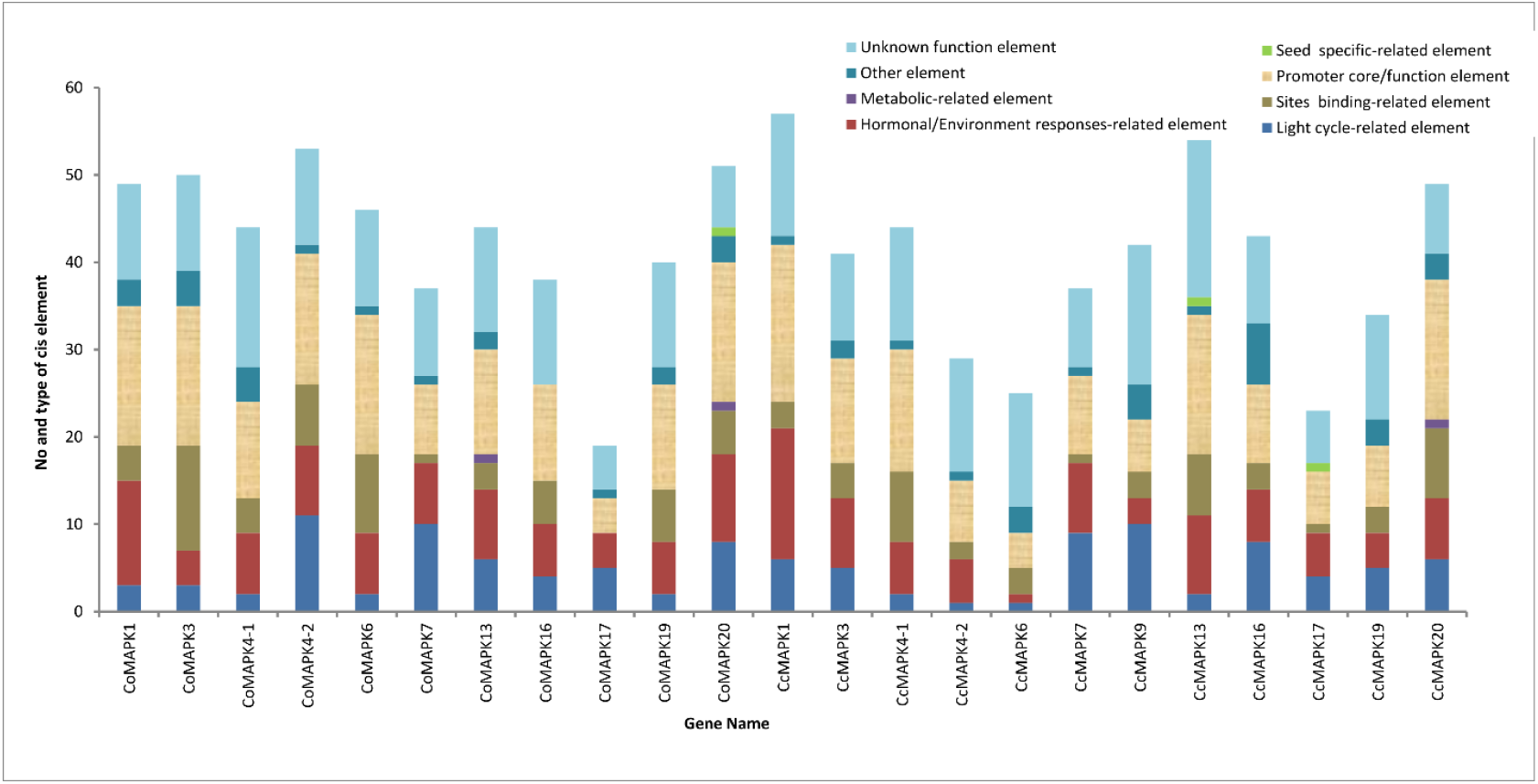
Promoter analysis of CoMAPK and CcMAPK genes. The numbers of different *cis*-elements were presented in the form of bar graphs; Different cis-elements with the same or similar functions are shown in the same color according to PlantCare promoter database.

A total of 218 cis-element found in the Co and CcMAPKs using PLACE database. Among them 41 elements are almost found in all the MAPKs (Fig. 6) and the most abundant CRE in all MAPK genes promoter is DOFCOREZM, which are specific DNA-binding proteins associated with the expression of multiple genes in plants. Besides this the abiotic stress related (ABRELATERD1, ACGTATERD1, CCAATBOX1, GT1GMSCAM4, MYB1AT, MYB2CONSENSUSAT), Ca^2+^ responsive (ABRERATCAL), light and circadian rhythms regulation (-10PEHVPSBD, IBOX, IBOXCORE, IBOXCORENT, SORLIP1AT, BOXIIPCCHS), phytohormone related (ARR1AT, ABREOSRAB21, ACGTABREMOTIFA2OSEM, ASF1MOTIFCAMV) and pathogen-related (GT1GMSCAM4, WBOXNTERF3) cis-elements in *MAPK* genes are also rich in both jute species (Fig. 6).

**Figure 6.**
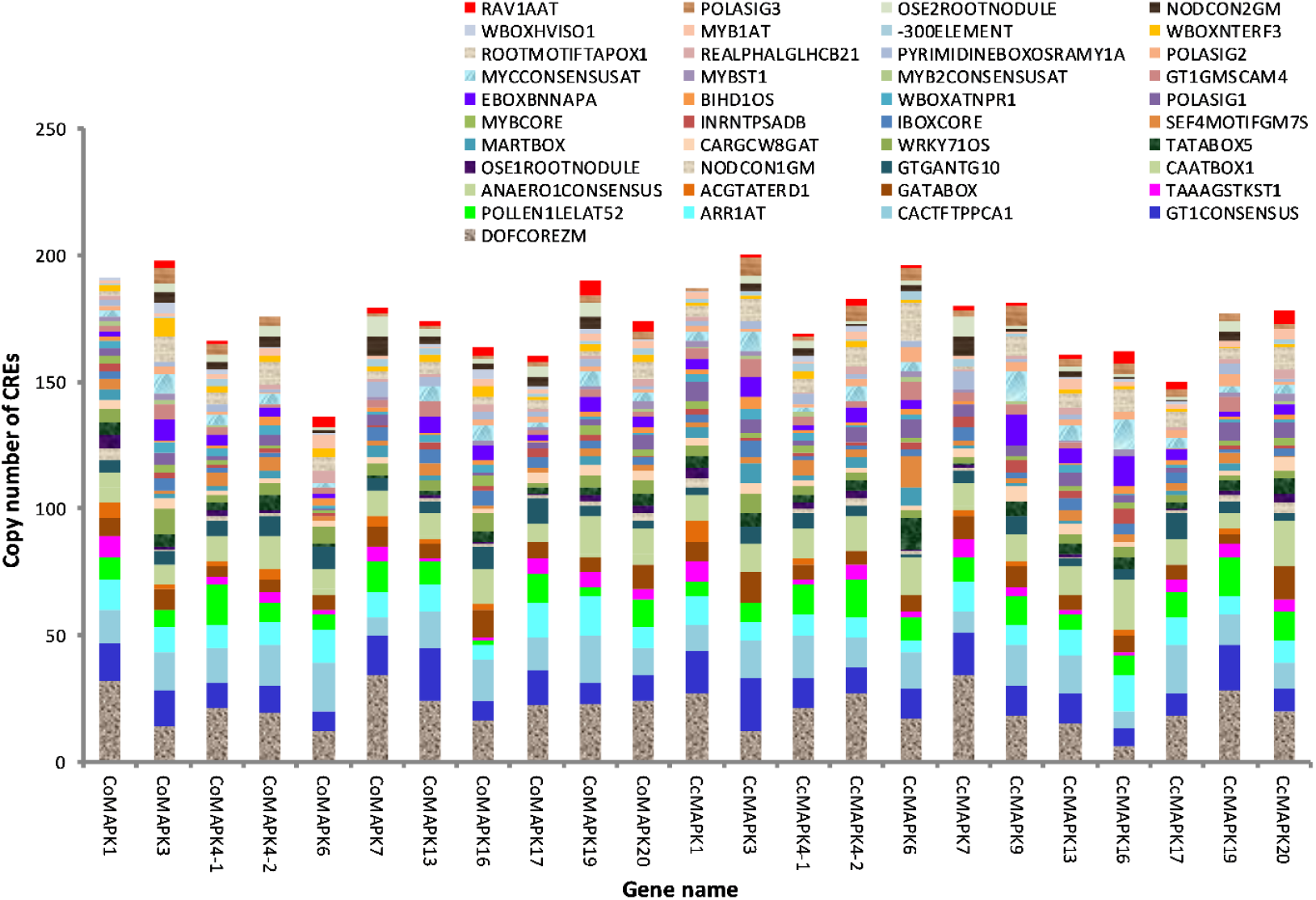
Promoter analysis of CoMAPK and CcMAPK genes. The numbers of different *cis*-elements were presented in the form of bar graphs; Different cis-elements with the same or similar functions are shown in the same color according to PLACE promoter database.

### Expression analysis of MAPKs during salt stress condition

Gene expression analysis can give a decisive idea about the function of a gene. To determine the role of *Co* and *CcMAPK* genes in response to adverse environment, we analyzed the available gene expression raw data (downloaded from NCBI) in root and leaf samples under salt and drought treatment. Moreover, we also analyzed the fiber vs seedling transcriptome data to delineate the role of MAPK signaling in fiber formation. Here we illustrate both *Corchorus* species MAPKs modification by salt and drought stress along-with bast fiber formation (Fig. 7). We investigated the expression pattern of MAPKs in leaves and roots treated with salt stress (Fig. 7a,). Exposure to salt upregulated the mRNA levels of six *CoMAPKS* (*CoMAPK1, CoMAPK3, CoMAPK4_1, CoMAPK4_2, CoMAPK6, CoMAPK7* and *CoMAPK19*) and six *CcMAPKs* (*CcMAPK3, CcMAPK4_1, CcMAPK6, CcMAPK7, CcMAPK9* and *CcMAPK16*) in both root and leaf in salt sensitive and salt tolerance genotype. On the other hand *CoMAPK16, CoMAPK20* and *CcMAPK1, CcMAPK17, CcMAPK20* shows down regulation in both leaf and root (Fig. 7a). Moreover, *CoMAPK17* and *CcMAPK13* had higher expression in root but lower expression in leaf where as *CcMAPK4_2* and *CcMAPK19* showed lower expression in root but expression is higher in leaf (Fig. 7a).

**Figure 7.**
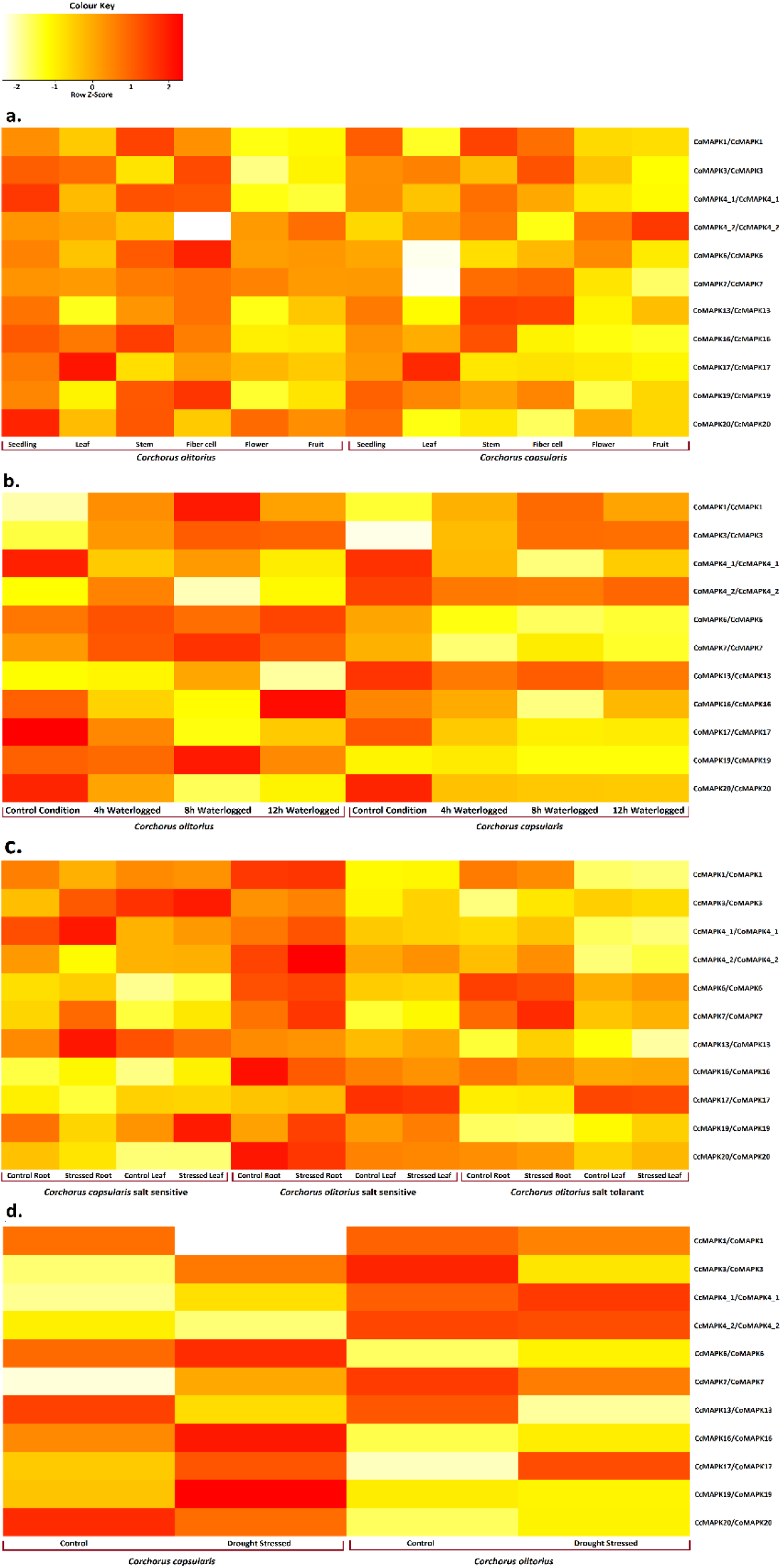
Expression profiling of Co and CcMAPK genes in six different tissues and against Waterlogged stress, Salt stress and drought stress. The log2 transformed FPKM (fragments per kilobase of exon model per million mapped reads) added by a pseudo-count of 1 value were used for heat map construction.

In drought stress, most *CoMAPK* and *CcMAPK* genes showed different expression profiles where four and seven MAPK genes were upregulated in *C. olitorius* and *C. capsularis* respectively. Moreover, *MAPK4.1, 6, 16, 17, 19* showed more expression in both species (Fig. 7b).

Furthermore, to examine the involvement of *MAPKs* in fiber cell development, we analyzed the expression of fiber cells against seedling. For the first time, this study reports that *MAPKs* have a potential role in fiber cell development. We found that seven and four *MAPKs* were highly expressed in fiber cell in *C. olitorius* (CoMAPK*1*, *3*, *4-1*, *7*, *13*, *16*, and *19*) and *C. capsularis* (*CcMAPK4-1*, *6*, *7* and *13*) respectively. The rest two *CoMAPKs* (*CoMAPK4-2* and *20*) and six *CcMAPKs* (*CcMAPK1, 4-2, 16, 17, 19* and *20*) expression is higher in jute seedlings. *CoMAPK6* and *17* and *CcMAPK3* has no expression in fiber cell (Fig. 7c).

## Discussion

In recent years, the gene families characterization has been suitable for studying their function(Tohge and Fernie, 2010). The accuracy and reliability of gene family characterization analysis depend on the genomic sequences. The availability of the decoded two *Corchorus* species genome sequence has made it possible to identify all the MAPK family members among these two species for the first time. Among the plant gene families Signaling gene of MAPKs plays an important role in multifaceted biological processes such as growth, development, and regulation of various environmental stresses in plants(Abass and Morris, 2013; Hamel et al., 2006; Jaggi et al., 2012; Jalmi and Sinha, 2015; Kim et al., 2016; Lee et al., 2009; Piao et al., 2018; Taj et al., 2010; Wang et al., 2017; Xu and Zhang, 2015). It is very important to assign an appropriate and specific name to each member of the family to enable a thorough understanding of it. The proper nomenclature of these MAP Kinase cascade genes should be used following an orthology or sequence homology-based MAPK gene nomenclature guidelines to maintain consistency across the plant kingdom. The accuracy and reliability of gene family characterization analysis depend on the genomic sequences. In this study, we identified 11 and 12 putative MAPK genes in the *C. olitorius* and *C. capsularis* genome respectively, which is slightly higher than papaya (9 members)(Mohanta et al., 2015), *Aquilegia coerulea* (10 members)(Mohanta et al., 2015), *Selaginella moellendorffii* (6 members)(Mohanta et al., 2015), *Physcomitrella patens* (8 members)(Mohanta et al., 2015) and similar to that of *Citrus sinensis* (12 members)(Mohanta et al., 2015), *Fragaria vesca* (11 members)(Zhou et al., 2017), Eucalyptus grandis (13 members)(Mohanta et al., 2015), *Ricinus communis* (12 members)(Mohanta et al., 2015), grapevine (14 members)(Cakir and Kilickaya, 2015) and strawberry (12 members)(Zhou et al., 2017), but smaller than in Arabidopsis (20 members)(Ichimura et al., 2002), *Solanum lycopersicum* (17 members)(Kong et al., 2012) *Helianthus annuus* (28 members)(Neupane et al., 2019) and apple (26 members)(Zhang et al., 2013). The full-length sequences of putative *Co* and *CcMAPKs* ranged from 368 to 612 amino acids. Variation in the length of the entire MAPK gene is usually due to differences in the length of the MAPK domain or the number of introns(Cakir and Kilickaya, 2015) (Fig.1 and 3). In the current study, the identified 11 and 12 *Co* and *CcMAPKs* were classified into four groups (A-D) according to their phylogenetic clades together with their MAPK orthology in Arabidopsis and Cacao which were consistent to the MAPKs previously identified in Arabidopsis(Colcombet and Hirt, 2008), poplar(Nicole et al., 2006), rice(Rao et al., 2010), Brachypodium distachyon(Chen et al., 2012), Malus domestica(Zhang et al., 2013), Ziziphus jujuba(Liu et al., 2017), Triticeae species(Goyal et al., 2018), Brassica rapa(Wu et al., 2017), and Fragaria vesca(Wei et al., 2017). Co and CcMAPK4-1, 4.2 and 13 belong to group A, which contain AtMAPK4, 5, 11, 12, 13 and TcMAPK4-1, 4-2 (Fig. 2). AtMPK4 is phosphorylated and activated by the upstream components AtMEKK1 and AtMKK2 upon cold and salt stress signaling in Arabidopsis (Ichimura et al., 2000; Teige et al., 2004). *Co* and *CcMAPK3* and *6* associate with group B, which includes *At* and *TcMAPK3* and *6* (Fig.2). It has been well-characterized that *AtMAPK3* and *6* is activated in response to biotic and abiotic stress(Ichimura et al., 2002; Jonak et al., 2002). The MAPKs group C contain Co and CcMAPK1 and 7 along with AtMAPK1, 2, 7 and 14 which are also activated by biotic and abiotic stress(Danquah et al., 2015). It is well studied that AtMAPK4 is crucial for activation of abiotic stress(Ichimura et al., 2001). Group D includes 4 and 5 *Co* and *CcMAPKs* respectively, which have the TDY motif in their Activation loop, which are consistently found in members of other MAPK groups. We found group D is the largest group in most plant species (Fig. 2).

The result of structural analysis of *Co* and *CcMAPK* was to some and extent, conserved according to phylogeney grouping indicating their evolutionary conservation (Fig.3). Interestingly, members in each group share the similar numbers of intron with corresponding groups of Arabidopsis, poplar, Tomato, Kiwi fruit, Sunflower and strawberry(Colcombet and Hirt, 2008; Kong et al., 2012; Neupane et al., 2019; Nicole et al., 2006; Wang et al., 2018; Wei et al., 2017). Concurrently, exon lengths of these species are clearly more conserved than intron length (Fig. 3). Similar gene structure between different species indicate the high conservation of these signaling component in the dicotylednous plant beyond the primary sequence identity.

Identified both *Corchorus* species MAPKs contain all canonical protein domains and motifs associated with MAPKs; fourteen MAPKs contain the TEY motif and nine MAPKs contain the TDY motif in the activation loop (Fig.1). A similar ratio between TEY MAPKs and TDY MAPKs was observed in other dicotyledonous species including *Fragaria vesca* (7 TEY MAPKs to 5 TDY MAPKs), Kiwifruit (14 TEY MAPKs to 4 TDY MAPKs), Cotton (36 TEY and 20 TDYMAPKs) and Cassava (13 TEY MAPKs to 8 TDY MAPKs)(Chen et al., 2020; Wang et al., 2018; Yan et al., 2016; Zhou et al., 2017). Furthermore conserved motifs present in *Co* and *CcMAPKs* were dissected using the MEME tool, which indicates motifs are conserved correspond to their phylogenetic grouping (Fig. 4). All the members of this family share similar eight conserved motifs (Motif 1, 2, 3, 4, 5, 6, 9, and 12) (Fig. 4). Motif 11, 14 and 15 are unique to group D, where as group B and C have the unique motif named Motif 10(Fig. 4). These results indicate that members that are close phylogenetically also possess similar conserved motifs and similarity also observed in Kiwi fruit, cassava, canola, B. rapa, and pomegranate (Liang et al., 2013; Lu et al., 2015; Ren et al., 2020; Wang et al., 2018; Yan et al., 2016).

GO analysis performed with Blast2Go describing the putative participation of *Co* and *CcMAPK* genes in multiple biological processes, molecular functions, and cellular component. The result of GO analysis demonstrate that most of the MAPKs are located in the nucleus, which is also supported by Plant-mLoc server and may involved in appropriate cellular adjustment in extracellular signals transduction(Sinha et al., 2011). In addition MAPK genes mainly involved MAPK cascade along with regulation of gene expression and abiotic stress responses and plant physiological process are occupied most of the proportion. The MAP kinase activity and ATP bindings are the most presented function in molecular function category. These features support the previous findings of MAPKs in barly(Cui et al., 2019).

The cis-regulatory elements in promoter region play crucial role for regulating gene expression. Therefore, the cis-regulatory elements of all *Co* and *CcMAPKs* were analyzed in two different platform one is PlantCare and another is PLACE (plant *cis* -acting regulatory DNA elements) database (Fig. 5, 6). In plantCare database the elements related to hormone and environment stress, photoreaction, promoter core element were notably abundant. Moreover, *Co* and *CcMAPK* promoters had abundant cis-elements associated with hormone, mainly ABRE (abscisic acid responsiveness), CGTCA-motifs and TGAC G-motifs (jasmonic acid responsiveness), indicating that regulation of MAPKs might be hormone dependent (Ding et al., 2016; Han et al., 2019) (Fig.5). The presence of stress-related cis-elements such as ABA-responsive cis-element (ABRE), stress response element (STRE), anaerobic responsiveness (ARE), low-temperature responsiveness (LTR), myeloblastosis (MYB) and its related promoter, MYB binding site (MBS), TC-rich repeats, wounding responsiveness (W-box) and pathogen responsiveness (WUN-motif), were found in the Co and CcMAPKs promoters like in other plant MAPKs promoter and indicating their gene expression during stress conditions(Kong et al., 2012; Liu et al., 2015; Singh et al., 2018-03) (Fig.5). Moreover we also analyzed *Co* and *CcMAPKs* in another promoter database Plant Cis-acting Regulatory DNA Elements (PLACE) where we found a lot of photosynthesis, plant hormone signaling, and organ development regulatory elements were accumulated, such as CACTFTPPCA1, DOFCOREZM, and ARR1AT (Fig.6). In addition, most portion of *cis*-acting DNA elements were related to biotic and abiotic stress response, especially to salt and drought stress, such as MYCCONSENSUSAT, EBOXBNNAPA, WRKY71OS, GT1GMSCAM4, CCAATBOX1, and ACGTATERD1 (Fig.6). Many of these cis-elements were also reported in the promoter of MAPKs of tomato (Kong et al., 2012), *B. distachyon* (Chen et al., 2012), kiwifruit(Wang et al., 2018) and cucumber (Wang et al., 2015), and they change their expression level on treatment with various stresses.

MAPK3, MAPK4 and MAPK6 have been reported to function in plant development and biotic/abiotic stress responses in Arabidopsis(Ichimura et al., 2002; Jonak et al., 2002; Kim et al., 2017; Sethi et al., 2014; Zhao and Zheng, 2017). Similarly increased expression of Co and Cc MAPK3, 4-1 and 6 indicates their involvement to salt tolerance. If we choosing the most effective single MAPK gene for both jute species for tolerant against salt and drought our findings reflect it as *Co* and *CcMAPK4-1*. There are various reports supporting the fact that overlapping signaling pathways operate in response to drought and salinity. In Arabidopsis, the MAPK pathway including MEKK1, MKK2, MPK4, and MPK6, is reported to be involved in salt, drought, and cold stress(Teige et al., 2004).

Furthermore, to examine the involvement of *MAPKs* in fiber cell development, we analyzed the expression of fiber cells against seedling. For the first time, this study reports that *MAPKs* have a potential role in fiber cell development. We found that seven *CoMAPKs* in from 11 and four from 12 CcMAPKs expression is higher in fiber cell. Among them CcMAPK7, 4-1 are highly expressed in both species, where as CcMAPK6 has two times more expression in fiber cell. Our finding support the previous studies in upland cotton(Chen et al., 2020).

## Conclusion

In this study, 11 and 12 putative MAPK genes in *C. olitorius* and *C. capsularis* were identified and investigated, and found to be comparable to those in other plants like Arabidopsis and Cacao. The MAPK family genes were compared and phylogenetically analyzed which allowed their classification into four groups, suggesting a high level of functional divergence. MAPK genes showed different temporal and spatial patterns of expression under salt stress conditions. Many of the MAPK genes were significantly upregulated by salt and drought stress conditions. The favored expression of MAPK genes provides a sound reason for their functional characterization in both *Corchorus* species. Furthermore, the fiber vs seedlings transcriptome result showed that MAPK signaling may plays a potential role when ultimate product of jute i.e. bast fiber is formed. It would be interesting to decipher the exact role of each of these MAPKs during growth, development, and abiotic stresses.

## Notes

### Competing Interest Statement

The authors have declared no competing interest.

